# Microbial pathway thermodynamics: structural models unveil anabolic and catabolic processes

**DOI:** 10.1101/2023.12.01.569601

**Authors:** Oliver Ebenhöh, Josha Ebeling, Ronja Meyer, Fabian Pohlkotte, Tim Nies

## Abstract

The biotechnological exploitation of microorganisms enables the use of metabolism for the production of economically valuable substances, such as drugs or food. It is, thus, unsurprising that the investigation of microbial metabolism and its regulation has been an active research field for many decades. As a result, several theories and techniques were developed that allow the prediction of metabolic fluxes and yields as biotechnologically relevant output parameters. One important approach is to derive macrochemical equations that describe the overall metabolic conversion of an organism and basically treat microbial metabolism as a black box. The opposite approach is to include all known metabolic reactions of an organism to assemble a genomescale metabolic model. Interestingly, both approaches are rather successful to characterise and predict the expected product yield. Over the years, especially macrochemical equations have been extensively characterised in terms of their thermodynamic properties. However, a common challenge when characterising microbial metabolism by a single equation is to split this equation into two, describing the two modes of metabolism, anabolism and catabolism. Here, we present strategies to systematically identify separate equations for anabolism and catabolism. Based on metabolic models, we systematically identify all theoretically possible catabolic routes and determine their thermodynamic efficiency. We then show how anabolic routes can be derived, and use these to approximate biomass yield. Finally, we challenge the view of metabolism as a linear energy converter, in which the free energy gradient of catabolism drives the anabolic reactions.

## Introduction

Microbial organisms are an essential part of most ecosystems. They function as vital members of natural production chains leading to the formation of chemical compounds that have complexity unreachable by current technological standards [29]. Not surprising, therefore, that much scientific effort has been spent to understand the metabolism of microbes necessary for the chemical interconversion of substances [34]. Today, the human exploitation of microbial metabolism has long left the stage of merely producing fermented products. Microbes, often viewed as natural factories, are used in biotechnological applications as an integrated component of drug and food fabrication or bioremediation projects [51, 41, 7]. However, the value of a microbial organism for economical usage depends on two main factors, metabolic capabilities, and thermodynamic constraints imposed on the microbial metabolism.

Gaining full knowledge about microbial metabolism was and is a complex scientific problem. In the pre-genomic era, researchers used precise measurements of input (substrate) and output (product) relationships for microbial cultures and a deep understanding of thermodynamics to design new biotechnological strategies, based on so-called macrochemical equations [32, 12, 13, 49, 34, 54]. A macrochemical equation summarizes the conversion of substrates into metabolic products and biomass. Describing microbial metabolism by a single macrochemical equation essentially treats microbial metabolism as a black box, ignoring all intracellular metabolic details. Still, this single chemical equation can accurately describe the overall metabolic activity and thus can serve to understand and predict biotechnologically important metabolic properties (see e.g. [12]).

The macrochemical equation can be understood as a sum of two separate reactions which describe catabolism (the breakdown of nutrients to gain free energy in the form of ATP) and anabolism (the formation of new biomass from the nutrients) [46]. In this picture, microbial growth is described as a thermodynamic energy converter, where the catabolic reactions provide the required free energy to drive anabolism (see Fig. 1). Here, the negative free energies of reaction of the catabolic and anabolic half-reactions, denoted by −∆_cat_*G* and −∆_ana_*G*, respectively, are generalised thermodynamic forces, and the respective reaction rates, *J*_cat_ and *J*_ana_, are generalised thermodynamic fluxes. The relation between these generalised forces and fluxes is often assumed to be linear [40, 54, 50, 47], following Onsager’s theory [25] for non-equilibrium thermodynamics. Onsager has shown that the linearity holds in general for systems close to equilibrium. Despite the attractiveness of the linear converter theory, it is not fully clear, to what extent this approximation is actually adequate for microbial growth. Regardless of these uncertainties, this simplified view of microbial growth as two coupled processes is insightful and allows estimating some principle thermodynamic limitations, such as maximally possible yields.

**Figure 1.**
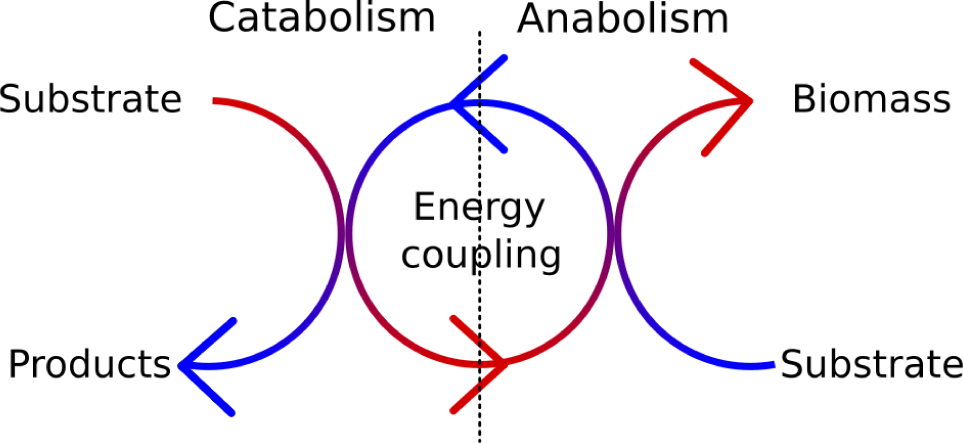
The view of microbial metabolism as a thermodynamic energy converter. Catabolic reactions have a large negative free energy gradient, driving anabolic reactions.

Describing macrochemical, catabolic and anabolic equations experimentally requires a precise measurements of all chemical substances that are consumed and produced by the growing microbes. These measurements are possible in controlled chemostat cultures, in which microbes grow on a defined growth medium. However, many mircobes cannot easily be cultured in chemically defined media, which prevents an experimental determination of macrochemical growth equations. However, with the recent scientific advancement in genome sequencing, genomic data became available for a huge number of organisms, including those which are difficult to culture [22]. This information greatly facilitates building a stoichiometric (or structural) metabolic models, manually or semi-automatically [11, 18, 20, 35]. With a structural model, metabolic capabilities can be systematically assessed and optimal flux distributions optimizing some objective function (such as biomass production) can be easily calculated [27]. Thus, such models can support strategies to improve the product yield in biotechnological applications. Elementary flux modes (EFM) are a systematic way to quantify the metabolic capabilities of an organism [36]. EFMs describe all possible pathways between substrate and products. However, due to combinatorial explosion [17], it is still challenging to calculate all EFMs for larger structural metabolic models, also with modern computational facilities. To overcome this, and because for many investigations only the conversion between substrate and product is relevant, elementary conversion modes (ECM) were introduced by Urbanczik and Wagner [44]. ECMs ignore all intracellular processes and only focus on the results of metabolic pathways. ECMs describe a minimal set of pathways that generates all steady-state substrate-to-product conversions [6]. Using ECMs instead of EFMs reduces the necessary computational power drastically. Additionally, modern software, such as `ecmtool` that allows parallelization of the computation, helps to obtain an exhaustive list of all metabolic capabilities of an organism in the form of ECMs [5].

As suggested in [6], we view ECMs as building blocks of macrochemical equations. We show how genomescale metabolic models can be used to systematically enumerate all possible catabolic pathways. With thermodynamic data, in particular energies of formation of substrates and products obtained from the `eQuilibrator` tool [2, 24], we characterise the catabolic pathways by their energy gradient. Using the network models, we further estimate the maximal ATP production capacity for each catabolic pathway, and thus determine their thermodynamic efficiencies.

We then analyze experimental data for *Saccharomyces cerevisiae* [31] and *Escherichia coli* [15], grown under controlled chemostat conditions in defined media, to separate the macrochemical equation into the catabolic and anabolic parts and characterise their thermodynamic properties. Interpreting our findings in the context of the energy converter model, we identify the limitations of the applicability of the linear converter model, but observe an interesting linear scaling law between growth rate and metabolic power.

## Results

### Calculating elementary conversion modes to characterise catabolic pathways

Genome-scale metabolic models are a formalisation of all known biochemical reactions of an organism. As such, they combine genomic, proteomic, and metabolic information to build an *in silico* representation that can be used to derive steady-state flux distributions. The inspection and analysis of genomescale metabolic models benefit from a rich theory for metabolic networks (see e.g. [37, 28, 43, 27, 16]). Here we use elementary conversion modes (ECMs, see [6, 5]) to assess the metabolic capabilities of several genome-scale metabolic networks. To illustrate our approach, we begin our analysis with the *E. coli* core network [26], an *E. coli* core metabolism model of reduced complexity, with only 72 metabolites connected by 95 reactions, of which 20 are exchange reactions. To systematically describe all theoretically possible catabolic routes, we used `ecmtool` [5] to calculate all ECMs which do not produce biomass. The resulting ECMs describe all possible routes and their stoichiometries by which external substances can be interconverted.

We then calculated for each individual ECM the standard Gibbs free energy of reaction, based on the standard energies of formation estimated by `eQuilibrator` [2]. The standard energies of catabolism ∆_cat_*G*^0^, normalised to one carbon mole of consumed substrate, are displayed in Fig. 2. Most of the catabolic pathways have relatively low energy gradients between approximately 20 and 50 kJ/Cmol. To this large group of pathways belong key catabolic routes, such as the fermentation of glucose to lactate or ethanol. Few catabolic routes exhibit Gibbs free energies of reactions with a higher energy gradient than 50 kJ/C-mol (around 17 ECMs). The pathways with the largest absolute Gibbs free energy of reaction are the combustion of glucose (∆_cat_*G*^0^ ≈ −488 kJ C-mol^−1^), and the production of formate from glucose (∆_cat_*G*^0^ ≈ −325 kJ C-mol^−1^). In particular, oxygen-using ECMs belong to the group with the largest absolute ∆_cat_*G*^0^ (compare red bars in Fig. 2). As shown in Fig. 2, usage of nitrogen does not appear to be an indicator of whether the respective ECM has a high or low energy gradient.

**Figure 2.**
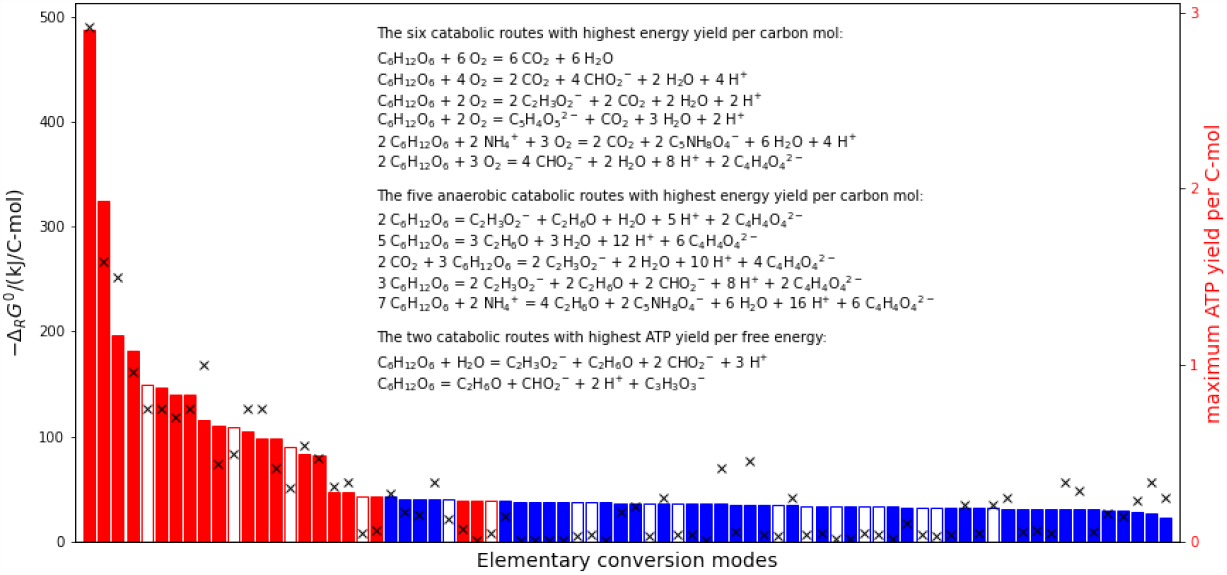
Standard Gibbs free energies of all catabolic pathways, normalised to carbon mole. The catabolic pathways were derived using elementary conversion modes (ECMs) calculated from the *E. coli* core network. Red symbolises ECMs that use oxygen, while blue denotes ECMs not using oxygen. Filled bars belong to ECMs that include no compounds with the element nitrogen while empty ones include nitrogen containing metabolites. The black crosses indicate the maximal yield of ATP per carbon mole nutrient for each ECM (right axis).

For each catabolic route, we use the metabolic model to calculate the maximal ATP yield. For this, the exchange reactions were constrained to the stoichiometries of the respective ECM, and subsequently the flux through ATP hydrolysis was maximized (see Methods). The resulting maximal ATP yields per carbon mole substrate are indicated by black crosses in Fig. 2. While as a tendency high energy gradient pathways also allow for a higher ATP yield, there are a considerable number of ECMs with very low ATP yield (34 ECMs exhibit a maximal ATP yield of less than 0.1 mol ATP per C-mol).

### Thermodynamic efficiency of catabolic routes

To investigate whether these general patterns are also conserved in more complete and therefore realistic genome-scale models, we repeated our analysis for the iJR904 metabolic network model [30] of *E. coli* as well as for the iND750 metabolic network model [9] of the yeast *S. cerevisiae*. Fig. 3 illustrates the results for *S. cerevisiae*, obtained with the iND750 model. We identified all ECMs using one of the four carbon sources glucose, xylose, *α*-ketoglutarate and pyruvate. Also here, most ECMs yield a low energy gradient, while those with the highest gradient correspond to the full oxidation of glucose and xylose. Full respiration for both sugars releases around 488 kJ C-mol^−1^ and are, thus, the ECMs with the highest free energy gradient. These two pathways also display the highest ATP yield per carbon mole (approx. 2.92 mol/Cmol) as indicated in Fig. 3, left panel. In contrast to this, most other catabolic routes release energy in a range between 40 and 200 kJ C-mol^−1^ with low ATP production. Among these routes are, besides other metabolic modes, fermentation reactions, such as the metabolisation of glucose to lactate or ethanol.

**Figure 3.**
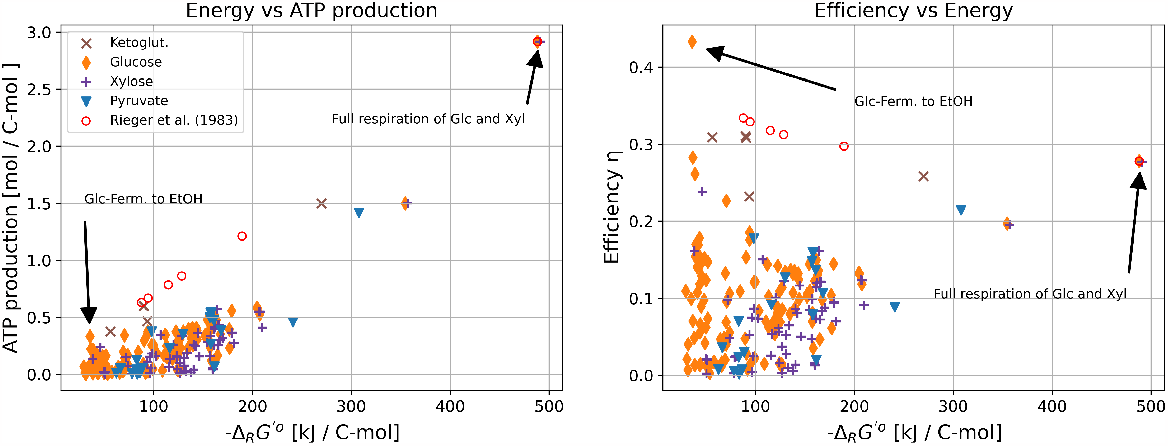
Thermodynamic characterisation of catabolic routes in *S. cerevisiae* genome-scale model (iND750) for *α*-ketoglutarate, glucose, xylose, and pyruvate as carbon source. Additionally, oxygen is allowed to be a substrate in the calculation of the elementary conversion modes. The efficiency is based on a typical value of 46.5 kJ/mol for production of ATP in *E. coli* [42].

The thermodynamic efficiency *η* of a thermodynamic engine is defined as the fraction of work generated per input energy provided to power the system. Defining the free energy used to drive ATP synthesis as the useful chemical work, we calculate the thermodynamic efficiency as

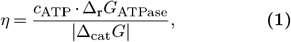

where *c*_ATP_ is the maximal ATP yield per carbon mol, ∆_r_*G*_ATPase_ the energy of reaction for ATP synthesis from ADP and inorganic phosphate, and ∆_cat_*G* the energy of reaction of the respective catabolic pathway. We assume the typical value of ∆_r_*G*_ATPase_ = 46.5 kJ mol^−1^ [42]. Further, we approximate the Gibbs free energy of catabolism by the corresponding standard Gibbs free energy of reaction, because changes in substrate and product concentrations in the medium are likely to have only a minor effect on the quantity. The determined efficiencies *η* are depicted in the right panel of Fig. 3. Interestingly, the pathways with the highest energy gradient are not the most efficient. For instance, under full respiration of glucose, only 28% of the released free energy is converted to chemical work producing ATP. In contrast, fermentation of glucose to lactate exhibits an efficiency of 43%. The fermentation of glucose to lactate is one of the most efficient reactions, with higher efficiencies only found in fermentation processes involving ethanol production.

### Deriving anabolic information from a metabolic network

Likewise, also anabolic pathways can be investigated in separation by employing genome-scale metabolic models. The theoretical limit how much carbon in the nutrient can be converted to biomass carbon is given by the degree of reduction. If biomass is more reduced than substrate, a fraction of the substrate carbons need to be oxidised in order to ensure the overall redox balance. Specifically, if *γ*_*S*_ and *γ*_*X*_ are the degrees of reduction of substrate and biomass, respectively, then the theoretical maximal yield is given by [32]

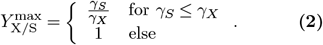

Based on the elemental composition of the biomass and the substrate, an ideal anabolic reaction stoichiometry can be determined. Assuming a substrate with a sum formula [S] = CH_*x*_O_*y*_ (normalised to Cmol) and biomass with [X] = CH_*a*_O_*b*_N_*c*_, *γ*_*S*_ = 4 + *x*− 2*y* and *γ*_*X*_ = 4 + *a*− 2*b*− 3*c* (assuming ammonia as nitrogen source, see [32]). If *γ*_*S*_ ≤ *γ*_*X*_ the stoichiometry reads

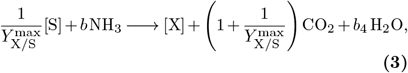

with 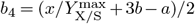 (see Methods).

This equation allows for the calculation of the standard energy of reaction of anabolism, ∆_ana_*G*^0^. To estimate the Gibbs free energy of formation of biomass, which is required to determine the energy of reaction, we employ the empirical method proposed by Battley [1].

Constraint-based models can be employed to investigate to what extent such an ideal anabolic reaction can be realised by a microorganism’s metabolism. We employ the genome-scale networks for *S. cerevisiae* and *E. coli* and minimise the nutrient uptake for a fixed biomass production, while allowing ATP to be provided externally (see Methods). A subsequent optimisation, in which the minimal nutrient uptake is fixed and the required reverse ATP hydrolysis is minimised, allows determining the minimal ATP requirement per carbon mole biomass formed. For the iJR904 model of *E. coli* metabolism, we obtain the following optimal anabolic stoichiometry for growth on glucose,

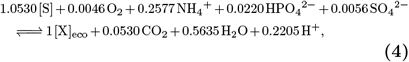

where [S] denotes 1 C-mol of substrate 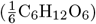 and [X]_eco_ 1 C-mol of *E. coli* biomass, with the sum formula determined by the biomass reaction of the iJR904 model

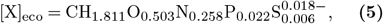

and a degree of reduction and energy of formation of

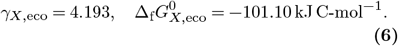

The calculated maximal yield of 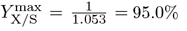 is slightly lower than expected by Eq. (2). This is explained by the fact that also small amounts of oxygen are required for the pure anabolic biomass formation. In iJR904 this is caused by a minimal required flux through a cytochrome oxidase which requires molecular oxygen as substrate.

A subsequent optimisation reveals a minimal requirement of 1.766 mol ATP per carbon biomass produced.

For the iND750 metabolic model of *S. cerevisiae*, we obtain

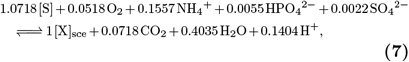

with

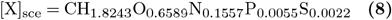

and a degree of reduction and energy of formation of

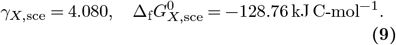

Here, the descrepancy between the computationally determined maximal yield of 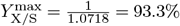 and the 98.0% expected from Eq. (2) is even larger.

The minimal requirement of ATP to produce biomass is predicted to be slightly larger than for *E. coli* with 2.031 mol ATP per C-mol biomass.

We repeated the calculations for different carbon sources. The results are summarized in Table 1. In general, the expected trend can be observed that the more oxidised carbon sources result in lower maximal yields. Moreover, the maximal yields predicted by the model are usually very close to the maximal yield predicted by the degree of reduction alone. Only for *S. cerevisiae* growing on oxoglutarate, yields predicted by the model are considerably lower. The reason for this is that the network defined by the iND750 model is not capable of producing biomass from oxoglutarate without metabolic side products. The optimal solution produces 0.073 mol xanthine (C_5_H_4_N_4_O_2_) per C-mol biomass. This leads to a reduced carbon (and in fact, nitrogen) yield, but a larger free energy gradient.

**Table 1.**
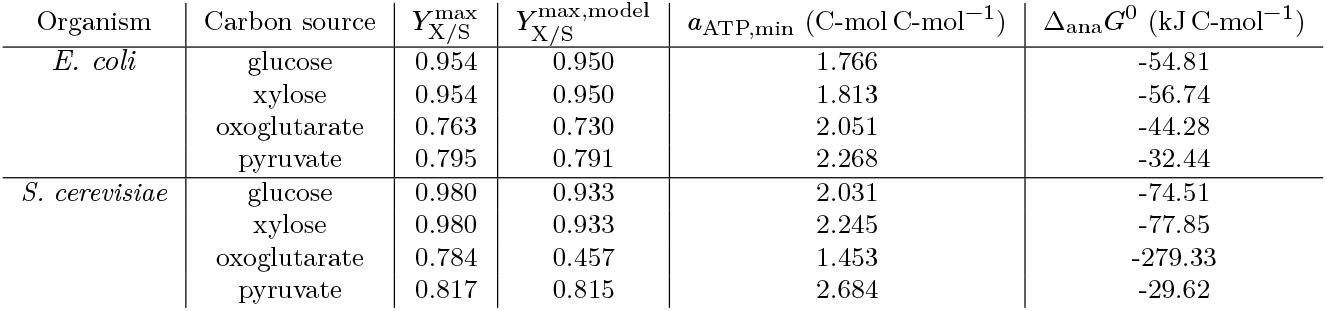
Thermodynamic properties of anabolic pathways. The theoretical maximal yield 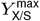 is calculated according to Eq. (2). The maximal yield 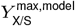 predicted by the metabolic model was determined using the linear program (22). The minimal anabolic ATP requirement per carbon mole biomass, *a*_ATP,min_ was determined using the linear program (23). The standard energies of reaction for anabolism, ∆_ana_ *G*^0^, were determined from overall anabolic stoichiometries like given for glucose in Eqs. (4) and (7).

| Organism | Carbon source | $Y_{X/S}^{\max}$ | $Y_{X/S}^{\max, \text{model}}$ | $a_{\text{ATP}, \min}$ (C-mol C-mol<sup>-1</sup>) | $\Delta_{\text{ana}}G^0$ (kJ C-mol<sup>-1</sup>) |
| --- | --- | --- | --- | --- | --- |
| *E. coli* | glucose | 0.954 | 0.950 | 1.766 | -54.81 |
|  | xylose | 0.954 | 0.950 | 1.813 | -56.74 |
|  | oxoglutarate | 0.763 | 0.730 | 2.051 | -44.28 |
|  | pyruvate | 0.795 | 0.791 | 2.268 | -32.44 |
| *S. cerevisiae* | glucose | 0.980 | 0.933 | 2.031 | -74.51 |
|  | xylose | 0.980 | 0.933 | 2.245 | -77.85 |
|  | oxoglutarate | 0.784 | 0.457 | 1.453 | -279.33 |
|  | pyruvate | 0.817 | 0.815 | 2.684 | -29.62 |

### Separating catabolism from anabolism based on chemostat data

In a controlled continuous microbial cultivation system, such as a chemostat [21, 14], it is possible to grow microbial cultures at a steady state with pre-defined growth rates. Measuring nutrient and gas exchange rates as well as nutrient and product concentrations in the reactor allows experimental determination of the overall growth stoichiometries [12, 13, 48, 49].

In the following we employ experimentally determined macrochemical equations for growth of *S. cerevisiae* [31] and *E. coli* [15] in chemostats at different dilution rates to calculate catabolic stoichiometries, ATP production potential, and thermodynamic efficiencies for each condition. The catabolic stoichiometry is calculated by first identifying the ideal anabolic stoichiometry based on the degrees of reduction of substrate and biomass, and then subtracting this anabolic stoichiometry from the macrochemical equation (for details, see Methods).

The determined catabolic coefficients are summarized in Fig. 4. It can clearly be seen that the onset of overflow metabolism, when glucose is partly fermented even in the presence of sufficient oxygen, occurs at growth rates of around 0.3 h^−1^ for *S. cerevisiae* and around 0.4 h^−1^ for *E. coli*. With the catabolic coefficients, we calculate the standard Gibbs free energy of reaction of the overall catabolic conversion, where we obtained the standard Gibbs free energies of formation, required for this calculation, from the `equilibrator` tool [2]. It can be observed (blue lines in Fig. 4) that with the onset of overflow metabolism, also the Gibbs free energy gradients are reduced significantly.

**Figure 4.**
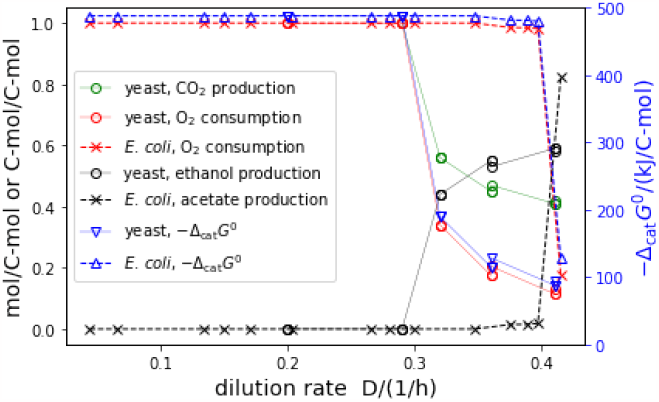
The catabolic stoichiometric coefficients and thermodynamic driving forces determined for chemostat growth of *E. coli* [31] and *S. cerevisiae* [15] at different dilution rates. All coefficients are given in mol/C-mol substrate, except for ethanol and acetate, which are given in C-mol/C-mol substrate. For *E. coli*, CO_2_ production is identical to O_2_ consumption. The thermodynamic driving force is given as standard energy of reaction of the overall catabolic conversion, normalised to one carbon mole of substrate.

### Is the linear energy converter a good model for microbial growth?

In the original publication that we draw our data from for *E. coli* [15], also higher dilution rates were investigated. For these, however, the carbon recovery rates were significantly below 90%, indicating that not all metabolic products were measured. The incomplete carbon recovery prevents a reliable calculation of catabolic stoichiometries. We therefore omitted these data, and will in the following focus on the energetic analysis of the metabolism of *S. cerevisiae*.

Microbial growth is often thermodynamically interpreted in the context of a linear energy converter model [50, 47], which assumes that the anabolic and catabolic fluxes linearly depend on the catabolic and anabolic forces, i. e. that

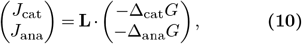

where the anabolic flux equals the growth rate, which in turn is set by the dilution rate of the chemostat, *J*_ana_ = *D*. In this model, the matrix **L** is the Onsager matrix of the phenomenolocigal coefficients [25].

Considering the large energy gradients, we approximate the actual Gibbs free energies by the standard energies. Moreover, we assume that the anabolic Gibbs free energy, ∆_ana_*G* is approximately constant over different dilution rates, because the stoichiometry of anabolism remains constant, according to Eq. (18). Based on the experimentally determined biomass composition of *S. cerevisiae* [31] (CH_1.79_O_0.57_N_0.15_), we calculate the standard energy of formation of biomass with the empirical method of Battley [1] to

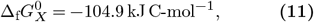

and with that the anabolic energy of reaction to

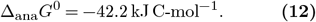

With this full knowledge of catabolic and anabolic fluxes and forces, we can challenge the linear converter model. In Fig. 5 (Fig. S2 for *E. coli*) various fluxes (catabolic glucose consumption, anabolic (growth) rate, total glucose consumption) are displayed as a function over the catabolic driving force or, alternatively (*x*-axis on top), the force ratio *x* = ∆_cat_*G/*∆_ana_*G*. It can clearly be observed that the fluxes do not linearly depend on the forces, as would be predicted by the linear converter model. On the contrary, fluxes are larger for smaller forces. This stark discrepancy from a linear converter model can readily be explained by considering that overflow metabolism results from active regulatory processes inside the cells. Often, overflow metabolism is explained by capacity constraints within the cell: whereas respiration results in a considerably higher yield, it also requires higher protein investment than fermentation, and therefore, at very high growth rates, it is more efficient to ‘waste’ substrate and operate a lower yield pathway (see e. g. [8, 23, 34]). This entails that there exist feedback regulation mechanisms, which are highly non-linear. It can be concluded that a linear energy converter model is too simplistic and is not in agreement with experimental data, at least over conditions in which the catabolic pathways change. The reason for this is to be sought in non-linear feedback mechanisms by which cells adapt their metabolism to external conditions.

**Figure 5.**
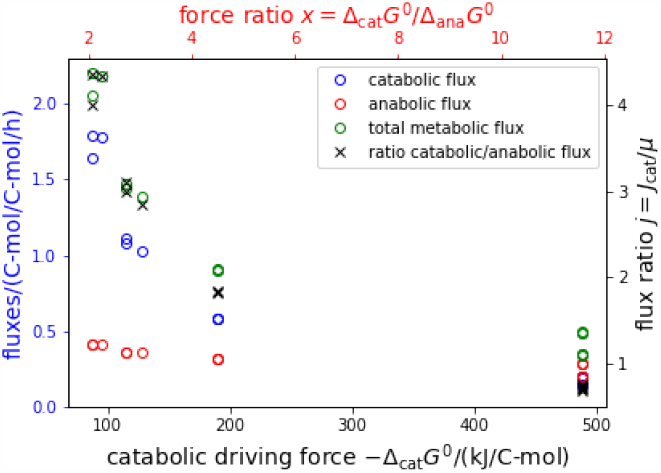
Metabolic fluxes as function of the catabolic driving force. Shown are the catabolic (blue), anabolic (red) and total (green) glucose consumption rates in dependence of the catabolic driving force, −∆_cat_ *G*^0^ . On the *x*-axis on the top, the force ratio *x* = ∆_cat_ *G*^0^ */*∆_ana_ *G*^0^ is given.

Similar to the calculation of maximal ATP yields of the different catabolic pathways, we determined the maximal ATP yield for the observed catabolic stoichiometries by constraining the genome-scale networks to the observerd catabolic stoichiometries. The red circles in Fig. 3 present the result of this calculation (left panel) and the respective thermodynamic efficiencies (right panel). Interestingly, the ATP yields as well as the efficiencies are higher than for elementary conversion modes with similar energy gradients. This can be explained by considering that the experimentally observed catabolic stoichiometries are a linear combination of two conversion modes (full respiration and fermentation) only, and that especially the fermentation pathways were identified to have the highest thermodynamic efficiency (see Figs. 2 and 3).

With the usual definition of the power as the product of flux and force, we can quantify the catabolic and anabolic powers *P*_cat_ = −*J*_cat_ *·* ∆_cat_*G* and *P*_ana_ = −*D ·* ∆_ana_*G*, as well as the power of ATP production *P*_ATP_ = *c*_ATP_ *· J*_cat_ *·* ∆_r_*G*_ATPase_ in kJ C-mol^−1^. Fig. 6 shows that the powers increase approximately linearly with growth rate, despite the fact that metabolism changes considerably for fast growth rates. Moreover, *E. coli* and *S. cerevisiae* behave rather similarly. For large growth rates, the powers of *E. coli* appear to be somewhat lower, but this could be a result from incomplete carbon recovery in the experiments, because besides acetate no other fermentation products were measured. This could lead to an underestimation of the Gibbs free energy gradients and consequently of the ATP yields.

**Figure 6.**
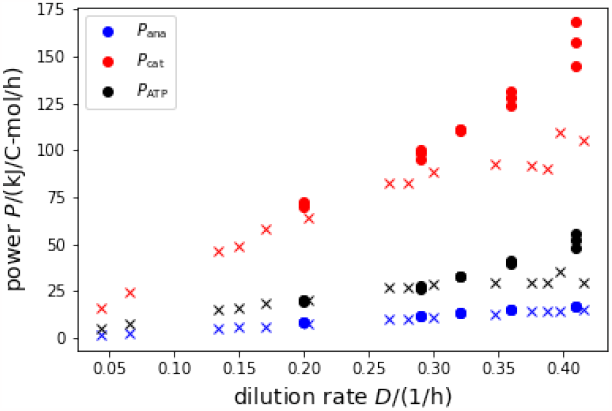
Catabolic and anabolic powers, as well as power of ATP synthesis as a function of growth rate. Catabolic power is depicted in blue, anabolic power in red, and ATP synthase power in black. Circles present results for *S. cerevisiae*, crosses for *E. coli*.

## Discussion & Conclusion

Microbial organisms are a cornerstone of the modern biotechnological industry. They are invaluable for producing pharmaceuticals, food, and construction materials [4]. Today, by using sophisticated genetic techniques and engineering, microorganisms can be used to tackle modern problems of society, such as the remediation of waste land, production of drugs or the finding environmental sustainable building materials [39, 45, 38]. However, such advancements in exploiting bacteria and unicellular eukaryotes were only possible with a thorough understanding of their metabolism. Multiple theories and techniques have been developed to gain knowledge about metabolic pathways. One of the currently most promising strategies is to develop genome-scale metabolic networks encoding almost all metabolic information available for an organism [10].

Genome-scale metabolic models are used to understand and probe the metabolic capabilities of an organism, and allow calculation of maximal yields [33]. The construction of such models only became possible with the advances in sequencing technologies, through which more and more fully sequenced genomes become available [22]. Before genome-scale metabolic networks, many scientists and engineers relied on macrochemical equations. These equations describe the metabolism of organisms as a black box by one overall chemical equation. Over the decades, this approach has been characterised extensively for its applicability and in the context of thermodynamics. However, a challenge when applying macrochemical equations is separating anabolism from catabolism so that both metabolic modes can be studied individually. Here, with the help of genome-scale models and thermodynamic calculations, we showed how we can extract both anabolic and catabolic conversions. We characterised both metabolic modes and challenged commonly used viewpoints on microbial metabolism, such as its representation as a linear energy converter.

By using elementary conversion modes (ECMs), which are an alternative to elementary flux modes, we could systematically enumerate catabolic routes from genome-scale networks of *E. coli* and *S. cerevisiae* (see Figs. S1 and 3). Combined with thermodynamic data of the Gibbs free energy of formation for all metabolites, as provided by the `eQuilibrator` tool, it is simple to derive standard Gibbs free energy of reactions for all input-output relationships (ECMs). By doing so, we calculated the catabolic driving force of microbial growth for all theoretically possible routes.

However, the second law of thermodynamics implies that not all of the available free energy can be used to perform useful chemical work. Combining ECMs with constraint-based modelling of genome-scale networks, we calculated the thermodynamic efficiency of the ATP production of each catabolic route. Interestingly, we find the most efficient pathways to exhibit a thermodynamic efficiency between approximately 30 and 45%. Interestingly, most catabolic routes show a considerably lower efficiency below 20%. It should be noted, though, that the values for the efficiency have to be interpreted with care. For one, we assumed standard energies of reaction for the catabolic routes and the actual concentrations of nutrients and catabolic products in the medium may slightly affect these values. Secondly, we have assumed a fixed value for the energy of reaction for ATP synthesis, which of course may change for different physiological conditions and depends primarily on the ATP:ADP ratio and the concentration of inorganic phosphate. Taking this into account, the highest efficiency, which is observed for the fermentation pathways, is very close to the 50% that is predicted to yield the highest ATP production rates by simple linear thermodynamic energy converter models [52]. It is remarkable that the actually realised catabolic pathways in chemostat cultures (see red circles in Fig. 3) provide a higher efficiency than elementary pathways with a similar free energy gradient. This observation stresses the important role of the pure respiration and fermentation pathways of catabolism. Because of their high efficiencies, operating them in combination always provides a higher thermodynamic efficiency than any single elementary conversion mode.

While a linear energy converter model seems adequate to predict optimal thermodynamic efficiencies of ATP producing pathways with a reasonable accuracy [52], our interpretation of experimental data shows that this is not the case when microbial growth is considered. Our results clearly demonstrate that the flux-force relationship is not linear, and that in fact anabolic and catabolic fluxes *decrease* with increasing catabolic driving force. In other words, the faster microbes grow, the lower the energy gradient that drives this growth. This observation, however, holds for conditions during which catabolism exhibits rather drastic changes, from pure respiration at low growth rates to largely fermentation at high growth rates. For growth in batch cultures on different substrate concentrations, however, it was shown that the linear converter model yielded very good results, which indeed fit the data better than a simple Monod equation [53]. It can therefore be hypothesized that the linear energy converter model is adequate as long as the catabolic mode does not change, and thus the driving force is mainly influenced by substrate concentration, but fails as too simplistic if experimental conditions encompass a change of catabolic pathways.

An interesting observation is that the output powers scale approximately linear with growth rate, and that the proportionality is very similar for two organisms as different as the bacterium *E. coli* and the eukaryote *S. cerevisiae*. For the anabolic power (growth rate times anabolic driving force), this result is trivial because we assumed the anabolic force, −∆_ana_*G*, to be constant. However, for the catabolic power (nutrient consumption rate for catabolism times catabolic driving force), this result is far from obvious. For technical systems, such as ships [3], bikes, cars or trains (see, [19] Chapter IIIA), the power increases over-proportionally with speed, approaching an approximately quadratic relationship. The linear power-growth rate relationship entails that, employing engineering terms, a “resistance” that needs to be overcome by the thermodynamic driving forces when producing new biomass is a constant rather than dependent on the biomass production rate. Moreover, the force, corresponding to the slope of the power-growth rate curves, appears to be the same for *E. coli* and *S. cerevisiae*. Whether these laws are of a universal nature remains to be tested with systematic experiments of more microbial species grown on different nutrient sources.

In summary, we could show how combining blackbox macrochemical approaches and genome-scale metabolic models can help to systematically characterise catabolic routes and find separate chemical equations for anabolism and catabolism. Interpreting experimental data from chemostats with our theoretical models reveals that the efficiency of catabolism appears optimal, both for *E. coli* and the yeast *S. cerevisiae*, over a wide range of growth rates. Moreover, our analyses allow us to speculate that the linear power-growth rate relationship is a universal property of microbial growth.

## Theory and Methods

### Calculating elementary conversion modes

Elementary conversion modes (ECMs, [44]) are a fast way to describe metabolic capabilities of an organism. We calculated ECMs to thermodynamically characterize all catabolic routes for three metabolic models, the *E. coli* core model [26], the genome-scale model iJR904 for *E. coli* str. K-12 substr. MG1655 [30], and the genome-scale model iND750 for the yeast *S. cerevisiae* [9], using `ecmtool` [6, 5]. For the genome-scale models, we hid all external metabolites that contain phosphate, sulfur, or nitrogen and dismissed all compounds with more than six carbon atoms. These steps reduced the number of the catabolic routes considerably and allowed the calculations to be performed in a reasonable time. For the two genome-scale networks, we focused as input (carbon source) for `ecmtool` on simple sugars and carboxylic acids (glucose, xylose, pyruvate, 2-oxoglutarate). Moreover, we allowed oxygen to be present. The output of `ecmtool` is a matrix, in which the rows are the respective elementary conversion modes, and the columns are all external metabolites that were not hidden. For the *E. coli* core network [26], no metabolite had to be hidden, and thus the full catabolic potential of the core network could be described.

### Estimating Gibbs free energy of reaction

To approximate the standard Gibbs free energy of reaction (∆_cat_*G*^0^) for each obtained ECM, we used the Python API of the `eQuilibrator` tool [24]. We extracted the Gibbs free energies of formation (∆_f_*G*^*?*^) for all external metabolites involved in an ECM. Next, we normalized the ECMs with respect to the carbon atoms of the carbon source (C-mol) and applied a Laplace transformation, adapting for temperature (298.15 K), pH (7.4), pMg (3.0), and ionic strength (0.25 M). We used Hess’s law to calculate the standard Gibbs free energy of reaction for each ECM,

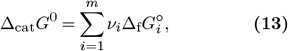

where *ν*_*i*_ and 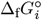 are the stoichometric coefficient and the Gibbs free energy of formation of the *i*^th^ external compound in the ECM, respectively.

For the calculation of the maximal ATP production for an ECM, we constrained all external fluxes to the values of the respective ECMs while maximizing ATP hydrolysis (excluding ATP maintenance):

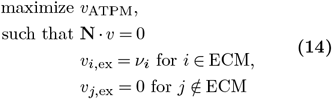

where **N** is the stoichiometric matrix of the metabolic model, *v*_ATPM_ is the flux through the reaction

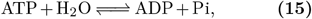

and *v*_*i*,ex_ are fluxes through the reaction exchanging metabolite *i*, which is constrained to the stoichiometric coefficient *ν*_*i*_ obtained by the respective ECM. The stoichiometric coefficients are normalised to one carbon mole substrate.

The thermodynamic efficiency of ATP production calculates as

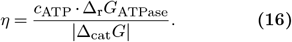

For the Gibbs free energy of ATP synthesis, we used a typical value for *E*.*coli* of ∆_r_*G*_ATPase_ = 46.5 kJ mol^−1^ [42].

### Calculating the stoichiometry of anabolism

We assume that the substrate [S] has the normalised sum formula CH_*x*_O_*y*_ and the biomass [X] has CH_*a*_O_*b*_N_*c*_, and that the biomass is more reduced than the substrate, i.e.

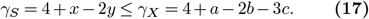

We assume an overall stoichiometry of

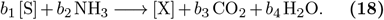

Every carbon that is converted to biomass will have to be reduced by *γ*_*X*_ −*γ*_*S*_ . From the overall redox balance it follows that for each carbon that is converted into biomass,

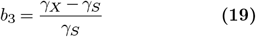

carbons have to be oxidised to CO_2_. From the carbon balance of (18) it follows that

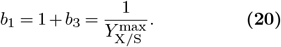

The nitrogen and hydrogen balances entail that

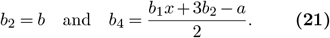

It is straight-forward to generalise these calculations to include sulfur and phosphorus into the biomass.

### Calcuating the optimal anabolic reaction

To determine the maximal yield and the minimal ATP requirement for biomass formation, we perform two subsequent linear programs. First, the exchange reactions are constrained, such that only the carbon source (substrate) and oxygen can be imported (negative flux), but other metabolites can be released (positive flux). The biomass reaction is constrained to one carbon mole per unit time. The ATP hydrolysis reaction is not constrained, which means it can run in reverse and provide ATP. Subsequently, substrate import (negative) is maximized:

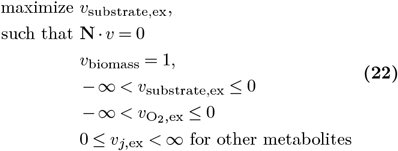

The resulting optimal flux is negative, and the absolute value denotes the minimal substrate requirement to produce one carbon mole of biomass, if ATP is provided in abundance.

In a second step, the determined minimal substrate requirement 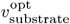 is fixed, and the ATP requirement is minimized by maximizing the (negative) flux through reaction (15):

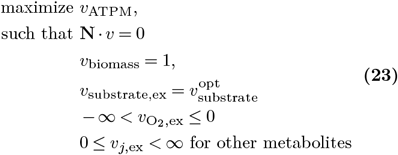

The absolute value of the optimal flux, |*v*_ATPM_| gives minimal ATP requirement for the production of one carbon mole biomass.

### Calculating the stoichiometry of catabolism

Macrochemical equations of the form

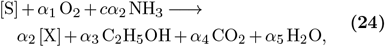

for *S. cerevisiae* (see [31]) and

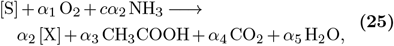

for *E. coli* were obtained from the original publications. Here, we use the notation [S] for one carbon mole of substrate and [X] for one carbon mole of biomass. The sum formula of biomass is assumed to be given by CH_*a*_O_*b*_N_*c*_ (hence the factor *c* in the stoichiometry of NH_3_), and is given in both cases in the original publication. The stoichiometric coefficients were obtained as follows. For *S. cerevisiae*, Table 1 in [31] already provides the stoichiometric coefficients for Eq. (24), which were, for our calculations, converted into carbon moles. For *E. coli*, we converted data from Table 2 in [15], which is given in g g^−1^ h^−1^ to C-mol C-mol^−1^ h^−1^, using the molecular weights of the chemical compounds as well as the biomass, normalised to one carbon mole.

The coefficients for the catabolic reaction

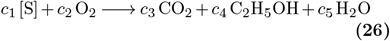

are simply determined by calculating (24) − *α*_2_ *·* (18), resulting in the coefficients

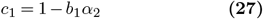

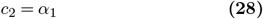

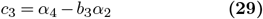

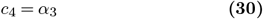

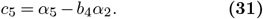

Subsequently, it is convenient to normalise this equation to the consumption of one carbon mole of substrate, i.e. dividing all coefficients by *c*_1_.

In the case of *E. coli*, acetate was excreted instead of ethanol at the onset of overflow metabolism [15]. In the calculation, ethanol can simply be replaced by acetic acid and the calculation remains identical.

## Abbreviations

EFM: elementary flux modes;
ECM: elementary conversion modes

## Funding

This work was funded by the Deutsche Forschungsgemeinschaft (DFG), project ID 391465903/GRK 2466 (T.N.) and the DFG by the Collaborative Research Center SFB1535, project ID 458090666/CRC1535/1 (F.P.). OE was supported by Deutsche Forschungsgemeinschaft under Germany’s Excellence Strategy – EXC-2048/1 – project ID 390686111.

## Author contributions

OE: initial idea and conceptualisation. OE: funding acquisition. OE, TN, FP, RM: visualisation. OE, TN, FP, RM, JE: formal analyses. OE, TN: writing—original draft and introduction. OE, TN, FP: writing—original draft and methods. OE, TN, FP: writing—original draft and results. OE, TN: writing—original draft, discussion, and OE, TN, FP, RM, JE writing—review and editing. All authors read and accepted the final version of the manuscript.

## Data Availability Statement

The original contributions presented in the study are included in the article/Supplementary Material, further inquiries can be directed to the corresponding author/s. The code can be found at https://gitlab.com/qtbhhu/thermodynamics-task-force/2023-energymetabolism-of-microorganisms.

## Conflict of interest

The authors declare that the research was conducted in the absence of any commercial or financial relationships that could be construed as a potential conflict of interest.

## Acknowledgements

The authors would like to thank Dr. St. Elmo Wilken for fruitful discussions and valuable input.

## Notes

### Competing Interest Statement

The authors have declared no competing interest.

