## Supplementary Material for "Microbial pathway thermodynamics: structural models unveil anabolic and catabolic processes"

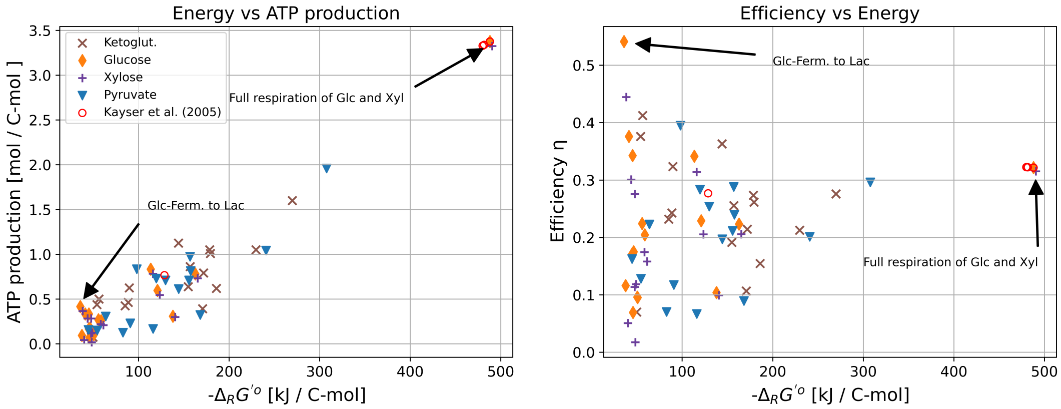


Figure S1: Thermodynamic characterisation of catabolic routes in *E. coli* genome-scale model (iJR904) for $\alpha$-ketoglutarate, glucose, xylose, and pyruvate as carbon source. Additionally, oxygen is allowed to be a substrate in the calculation of the elementary conversion modes. The efficiency is based on a typical value of 46.5 kJ/mol for production of ATP in *E. coli* [1].

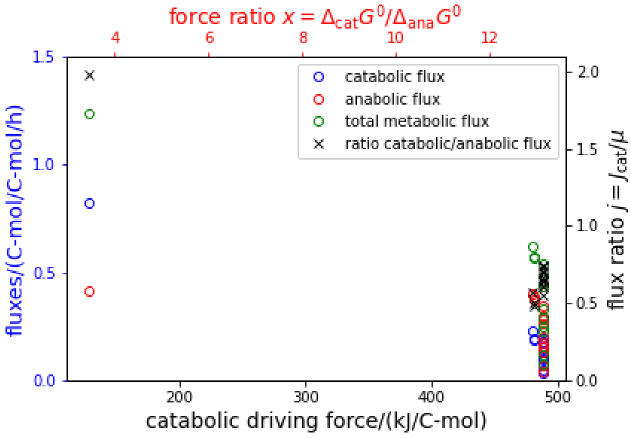


Figure S2: Metabolic fluxes as function of the catabolic driving force for data for *Escherichia coli* [2]. Shown are the catabolic (blue), anabolic (red) and total (green) glucose consumption rates in dependence of the catabolic driving force, $-\Delta_{cat}G^0$. On the $x$-axis on the top, the force ratio $x = \Delta_{cat}G^0/\Delta_{ana}G^0$ is given.
